# BART: a transcription factor prediction tool with query gene sets or epigenomic profiles

**DOI:** 10.1101/280982

**Authors:** Zhenjia Wang, Mete Civelek, Clint L. Miller, Nathan C. Sheffield, Michael J. Guertin, Chongzhi Zang

## Abstract

**Summary:** Identification of functional transcription factors that regulate a given gene set is an important problem in gene regulation studies. Conventional approaches for identifying transcription factors, such as DNA sequence motif analysis, are unable to predict functional binding of specific factors and not sensitive to detect factors binding at distal enhancers. Here we present Binding Analysis for Regulation of Transcription (BART), a novel computational method and software package for predicting functional transcription factors that regulate a query gene set or associate with a query genomic profile, based on more than 6,000 existing ChIP-seq datasets for over 400 factors in human or mouse. This method demonstrates the advantage of utilizing publicly available data for functional genomics research.

**Availability:** BART is implemented in Python and available at http://faculty.virginia.edu/zanglab/bart

## INTRODUCTION

Transcriptional regulation of gene expression plays a critical role in many cellular processes, including cancer development and progression (Bradner *et al.*, 2017; Lambert *et al.*, 2018). Identification of functional transcription factors is essential for understanding gene regulatory mechanisms in such processes. In gene expression profiling studies, ontology and pathway analyses (Huang *et al.*, 2008; McLean *et al.*, 2010; Subramanian *et al.*, 2005) can identify functional annotations of differentially expressed genes; however, this approach is unable to predict transcription factors that regulate those gene sets. Most existing methods for cis-regulatory prediction rely upon detecting overrepresented DNA sequence motifs near the gene promoters to identify sequence-specific DNA-binding factors (Boeva, 2016). Such methods are limited by the context-specific nature of transcription factor activity and by multiple factors sharing similar motifs (Jolma *et al.*, 2013). Moreover, most cis-regulatory events in mammalian genomes occur at distal enhancers, which cover much larger regions than promoters but without direct assignment to target genes; these regions are usually difficult to capture by motif scan alone (Shlyueva *et al.*, 2014).

Several methods have been developed to overcome these limitations of motif-based, promoter-biased approaches by using comprehensive epigenomic information (Dozmorov, 2017), such as DNaseI hypersensitive sites (Sheffield *et al.*, 2013). Model-based Analysis of Regulation of Gene Expression (MARGE) is a method developed for modeling differential gene expression using a compendium of public H3K27ac ChIP-seq datasets (Wang *et al.*, 2016). By quantifying the regulatory potential of active enhancer histone mark H3K27ac on each gene in the genome from each ChIP-seq dataset, MARGE uses a semi-supervised learning approach to predict a genome-wide cis-regulatory profile for any query gene set. Leveraging over 6,000 transcription factor ChIP-seq datasets available in the public domain (Mei *et al.*, 2017), we have developed Binding Analysis for Regulation of Transcription (BART), a new method for prediction of functional transcription factors by associating ChIP-seq binding information with MARGE-predicted genomic cis-regulatory regions.

## METHODS

BART identifies transcription factors whose genomic binding profile correlates with a query cis-regulatory profile derived from either a gene set or a ChIP-seq dataset (Fig. 1). BART uses a previously curated union DNaseI hypersensitive sites (UDHS) as a repertoire of all cis-regulatory regions in the genome (Wang *et al.*, 2016). The UDHS has over 2.7 million sites in human or 1.5 million in mouse, covering regions of 307 Mbp or 121 Mbp in the human or mouse genomes, respectively, including most enhancers and promoters. We used UDHS to represent genomic cis-regulatory regions, because for the vast majority of transcription factor ChIP-seq datasets, over 80% of the identified peaks (binding sites) are overlapped with UDHS (Suppl. Fig. S1).

**Fig. 1.**
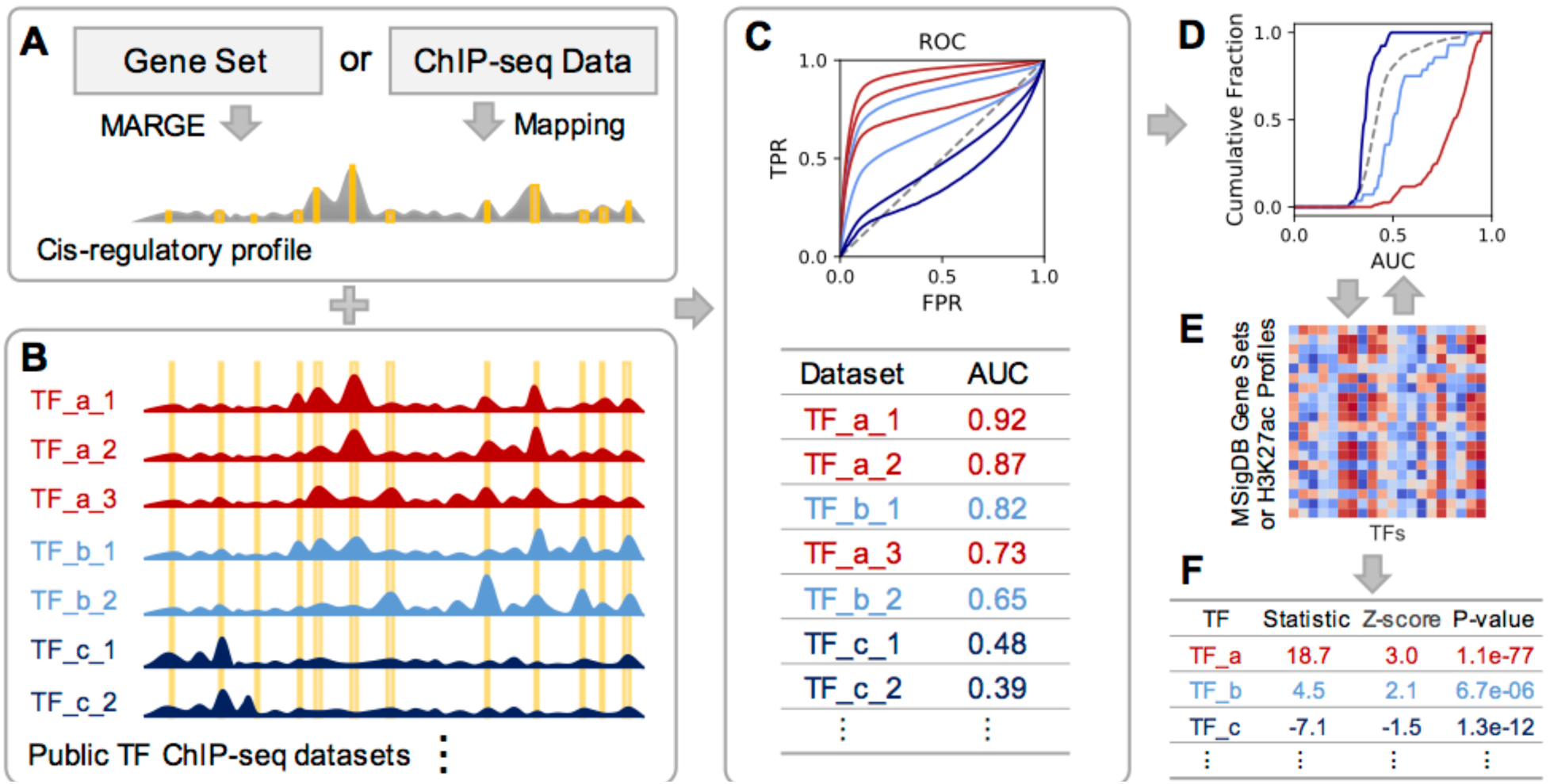
BART workflow. (A) Cis-regulatory profile is generated from query gene set by MARGE or from a ChIP-seq dataset by genomic mapping. Yellow bars indicate UDHS. (B) Each transcription factor binding profile from a ChIP-seq dataset is converted to a binary string showing presence of absence at each UDHS. (C) Top: Each ROC curve represents the prediction performance of a transcription factor profile from B by the query cis-regulatory profile from A; Bottom: Area under the ROC curve (AUC) is calculated for all datasets. (D) AUC are grouped by factor, and Wilcoxon test is performed for each factor compared with all datasets as background. In this example, cumulative distributions show significantly higher AUC for TF_a (red). (E) Wilcoxon test statistic is calculated for each transcription factor from each dataset in the background for Z-score calculation. (F) BART outputs a ranked list of all transcription factors.

Given a query gene set as input (“*geneset*” mode), MARGE is applied to predict a genomic cis-regulatory profile that regulates the gene set. For a ChIP-seq dataset input (“*profile*” mode), BART will use the ChIP-seq read count near each UDHS site to quantify the UDHS and generate the cis-regulatory profile (Fig. 1A).

For each transcription factor ChIP-seq dataset from the collected data compendium (Mei *et al.*, 2017), we map the identified peaks to the UDHS and assign each UDHS site a binary score indicating whether this site overlaps with a peak (Fig. 1B). To assess whether the query cis-regulatory profile (as a ranking of the UDHS) is associated with each ChIP-seq profile, we generate a receiver operating characteristic (ROC) curve using the factor-bound UDHS as true positive and the cis-regulatory rank as the predictor. We use the area under the ROC curve (AUC) score to quantify the association between the cis-regulatory profile and each dataset (Fig. 1C).

We then apply statistical tests to assess the significance of each factor using the AUC scores from all transcription factor ChIP-seq datasets. Given that each factor can have multiple ChIP-seq datasets in the data compendium generated under different biological conditions or by different labs, we first use the Wilcoxon rank-sum test for each factor by comparing the AUC scores from ChIP-seq datasets of that factor with the AUC scores from all datasets (Fig. 1D). Here we take advantage of the collective information from many ChIP-seq datasets for one factor as a trade-off of the cell-type specificity of transcription factor binding patterns, considering that the genomic binding profiles of the same factor across different cell types are usually more similar than the binding profiles of different factors in the same cell type (Suppl. Fig. S2). To assess the specificity of each factor, we build up background models using the Wilcoxon test statistics obtained from each annotated gene set from the Molecular Signatures Database (MSigDB) (Liberzon *et al.*, 2011) in the *geneset* mode, or using each H3K27ac ChIP-seq dataset from the ChIP-seq data compendium (Wang *et al.*, 2016) in the *profile* mode, respectively (Fig. 1E, Suppl. Fig. S3). We then calculate a standard Z-score for each factor, by comparing the test statistic from the query data with those from the background. Finally, we obtain the transcription factor prediction based on the average rank of Wilcoxon P-value, Z-score, and the maximum AUC among datasets for that factor (Fig. 1F). More details can be found in Supplementary Data.

## RESULTS AND DISCUSSION

We tested BART on several gene sets derived from differentially expressed genes after activation or inhibition of known transcription factors, including ESR1, AR, NR3C1, PPARG, NOTCH1, and POU5F1 (Wang *et al.*, 2016). In the BART result, the true functional factor was ranked on top (1/454) of the candidates in 4/6 gene sets; and ranked No.2 and No.47 for ESR1 and NR3C1, respectively (Suppl. Fig. S4). The highest ranked factor predicted from NR3C1 target genes is NR2A1, another nuclear receptor. The correct predictions are robust and not affected by randomness in MARGE outputs (Suppl. Fig. S5). These results indicate that BART can successfully predict transcription factors that regulate a given gene set. To evaluate the performance of BART, we compare BART with 4 other transcription factor prediction tools that take a gene set as query, including the ENCODE ChIP-seq Significance Tool (Auerbach *et al.*, 2013), HOMER (Heinz *et al.*, 2010), iRegulon (Janky *et al.*, 2014), and Pscan (Zambelli *et al.*, 2009) (Suppl. Table 1). On prediction of the true factor from the 6 gene sets, BART outperforms other methods for 5 cases, except NR3C1 (Suppl. Table 2).

BART can identify transcription factors that regulate any gene set or associate with any genomic profile. BART provides functional interpretations to differential gene expression analysis. BART makes predictions based on direct binding information from public ChIP-seq data only, as an orthogonal approach to conventional DNA sequence motif search. It focuses on transcription factor binding at open chromatin regions in the genome represented by UDHS, most of which are located at enhancers far from coding genes, providing an effective and extensive coverage for cis-regulatory events in the genome.

BART can have a broad application in transcriptional regulation studies. Leveraging abundant information from public data, BART can help generate hypotheses about functional transcriptional regulatory mechanisms in any human or mouse cell system, especially the cases where functional transcription factors are unknown, such as effects of external treatments like drugs or exposures, during a developmental process, and comparing tumor with normal cells. Any transcriptomic profiling experiment that generates differentially expressed gene sets comparing two states can be subject to BART analysis. It can also be applied to validate the knock-down or knock-out experiments and help look for novel co-factors associated with known transcription factors.

When applying BART for transcription factor prediction, users should also be aware of several limitations of this method. First, BART prediction is based on information from existing ChIP-seq data. Even though we have included as many ChIP-seq datasets as possible and will continue to update the compendium as more data become available, there are still many factors that do not have publicly available ChIP-seq data in any cellular system. Second, in the “*geneset*” mode, the cis-regulatory profile prediction using MARGE is based on H3K27ac signals. Although H3K27ac has been shown as the histone mark that separates active enhancers from poised enhancers (Creyghton *et al.*, 2010; Rada-Iglesias *et al.*, 2012), other histone modifications such as H3K4 methylations have also been implicated as hallmarks of enhancers (Henriques *et al.*, 2018). Similar approaches can be adopted on other histone marks alone or collectively as extension of MARGE and BART in the future. Third, while taking advantage of the collectivity of multiple binding profiles of the same transcription factor in different cell types and conditions, BART does not particularly focus on the cell-type specificity of transcription factor binding patterns. The considerations are: 1) Cell-type specificity of the query gene set has been addressed in the MARGE prediction, where the regression-selected H3K27ac profiles contain the cell/tissue type information (Wang *et al.*, 2016); 2) The genomic binding patterns of the same factor across different cell types are more similar than the binding patterns of different factors in the same cell type (Suppl. Fig. S2). Therefore, in case the true transcription factor’s ChIP-seq data in the right cell type is not available, BART would still be able to identify the true factor from its similar binding patterns in different cell types, rather than finding a false factor from that same cell type. After all, due to the incomplete coverage of cell and tissue types from public chromatin accessibility profiling data and ChIP-seq data, the ability of BART in identifying transcription factors binding at specific cis-regulatory regions in an uncharacterized cell system is limited. In those cases, we suggest using BART in combination with other approaches such as motif- or co-expression-based methods to improve predictions. Despite these limitations, BART provides a framework for accurate prediction of functional transcriptional regulation by utilizing public data.

